# Asexual but not clonal: evolutionary processes in populations with automictic reproduction

**DOI:** 10.1101/081547

**Authors:** Jan Engelstädter

**Author notes:** Address: The University of Queensland, School of Biological Sciences, Brisbane, QLD 4072, Australia.

## Abstract

Many parthenogenetically reproducing animals produce offspring not clonally but through different mechanisms collectively referred to as automixis. Here, meiosis proceeds normally but is followed by the fusion of meiotic products that restores diploidy. This mechanism typically leads to a reduction in heterozygosity among the offspring compared to the mother. Following a derivation of the rate at which heterozygosity is lost at one and two loci, depending on the number of crossovers between loci and centromere, a number of models are developed to gain a better understanding of basic evolutionary processes in automictic populations. Analytical results are obtained for the expected equilibrium neutral genetic diversity, mutation-selection balance, selection with overdominance, the rate of spread of beneficial mutations, and selection on crossover rates. These results are complemented by numerical investigations elucidating how associative overdominance (two off-phase deleterious mutations at linked loci behaving like an overdominant locus) can in some cases maintain heterozygosity for prolonged times, and how clonal interference affects adaptation in automictic populations. These results suggest that although automictic populations are expected to suffer from the lack of gene shuffling with other individuals, they are nevertheless in some respects superior to both clonal and outbreeding sexual populations in the way they respond to beneficial and deleterious mutations. Implications for related genetic systems such as intratetrad mating, clonal reproduction, selfing as well as different forms of mixed sexual and automictic reproduction are discussed.

## Introduction

The vast majority of animals and plants reproduce via the familiar mechanism of sex (Bell 1982): haploid gametes are produced through meiosis, and these fuse to form diploid offspring that are a genetic mix of their parents. Conversely, bacteria, many unicellular and some multicellular eukaryotes reproduce clonally, i.e. their offspring are genetically identical to their mother. These two extreme genetic systems can also be alternated, e.g. a few generations of clonal reproduction followed by one round of sexual reproduction. Such systems are found in many fungi (e.g., yeast) but also in animals such as aphids that exhibit ‘cyclical parthenogenesis’. However, there are also genetic systems that resist an easy classification into ‘sexual’ and ‘asexual’. Among them are automixis and related systems in which a modified meiosis takes place in females, leading to offspring that develop from unfertilized but diploid eggs and that may be genetically diverse and distinct from their mother (Stenberg and Saura 2009; Suomalainen *et al.* 1987). Explicably, there is much confusion and controversy about terminology in such systems, with some authors referring to them as asexual (because there is no genetic mixing between different lineages) and others as sexual (e.g., because they involve a form of meiosis and/or resemble selfing). Without entering this debate, I will adopt the former convention here, acknowledging that the latter is also valid and useful in some contexts. Also note that clonal, ‘ameiotic’ reproduction in animals is usually referred to as apomixis but that this term has a different meaning in plants (Asker and Jerling 1992; van Dijk 2009).

A good starting point for understanding automixis is to consider a specific system, and one that is particularly well-studied is the South African honeybee subspecies *Apis mellifera capensis,* the Cape honeybee (reviewed in Goudie and Oldroyd 2014). Within *A. m. capensis,* workers can lay unfertilized eggs that develop parthenogenetically into diploid female offspring via a mechanism called ‘central fusion’ (Suomalainen *et al.* 1987; Verma and Ruttner 1983). Here, meiosis occurs normally producing four haploid nuclei, but diploidy is then restored through fusion of the egg pronucleus with the polar body separated in meiosis I. This means that in the absence of recombination between a given locus and its associated centromere, the maternal allelic state at this locus is restored and in particular, heterozygosity is maintained. However, crossover events between a locus and its centromere can erode maternal heterozygosity, leading to offspring that are homozygous for one allele (see below for details and Fig.1). Although workers can produce diploid female offspring asexually, queens (which may be the daughters of workers) still mate and reproduce sexually. However, this system has also given rise to at least three lineages (two historical and one contemporaneous) that reproduce exclusively through central fusion automixis and parasitize colonies of another, sexual honeybee subspecies (*A. m. scutellata*). The contemporaneous lineage (colloquially referred to as the “Clone” in the literature) appeared in 1990 and has been spreading rapidly since, causing the collapse of commercial *A. m. scutellata* colonies in South Africa (the ‘Capensis Calamity’: Allsopp 1992). Heterozygosity levels are surprisingly high in this lineage given its mode of automictic reproduction (Baudry *et al.* 2004; Oldroyd *et al.* 2011). Initially, it was hypothesized that this is due to suppression of recombination (Baudry *et al.* 2004; Moritz and Haberl 1994), as this would make central fusion automixis akin to clonal reproduction. However, more recent work indicates that it is more likely that natural selection actively maintains heterozygosity (Goudie *et al.* 2012; Goudie *et al.* 2014).

**Figure 1.**
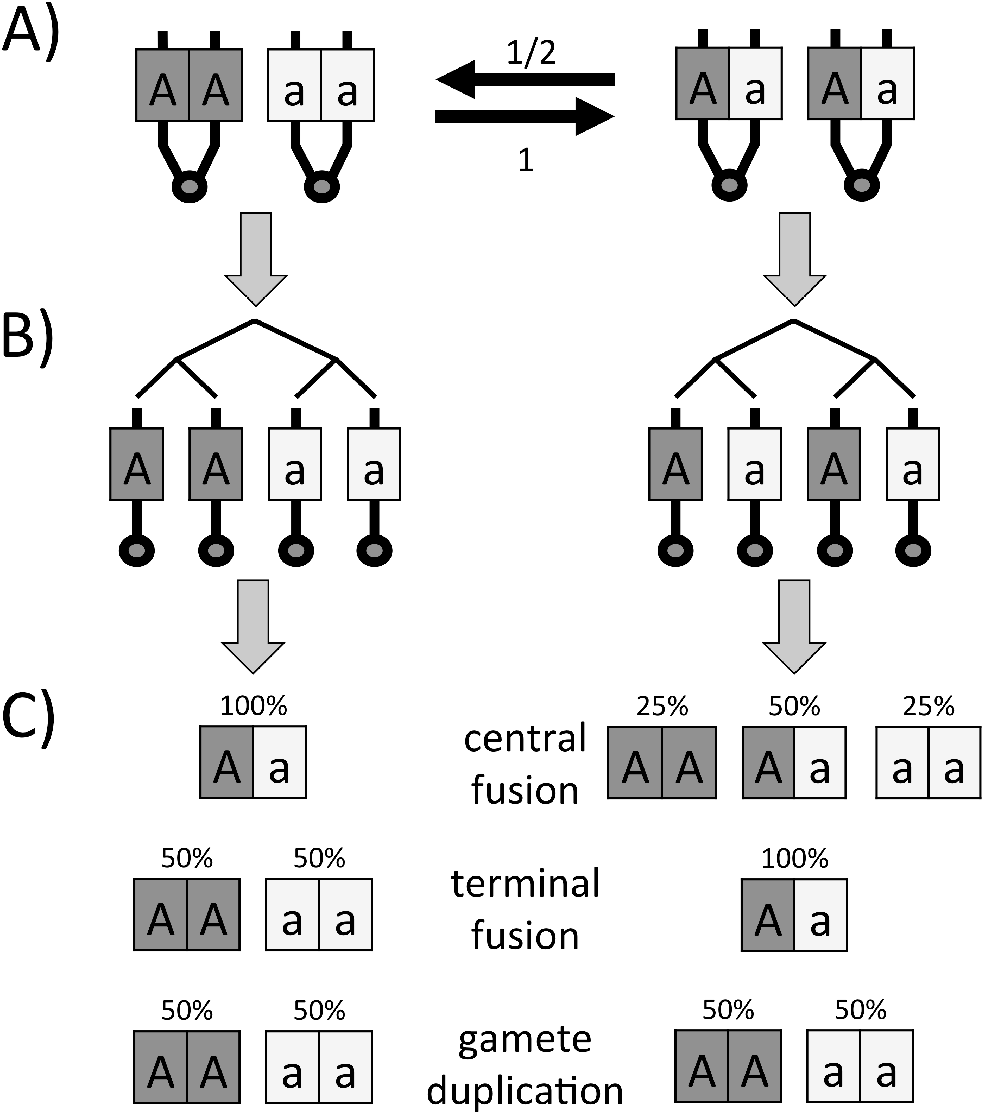
Illustration of the genetic consequences of automixis at a single locus. A) Starting from a heterozygous mother, during Prophase I, crossover events between the locus can induce switches between two possible states. Each crossover converts the original state where identical alleles are linked to the same centromere to the opposite state where different alleles are linked to one centromere. With probability ½, crossovers can then revert the second to the first state. B) Depending on which state is reached, meiosis will result in one of two possible genetic configurations. C) Fusion of meiotic products or suppression of the first mitotic division can then lead to different proportions of zygote genotypes.

Several other species also reproduce exclusively or facultatively through central fusion automixis, including other hymenopterans (e.g., Belshaw and Quicke 2003; Beukeboom and Pijnacker 2000; Oxley *et al.* 2014; Pearcy *et al.* 2006; Rabeling and Kronauer 2013), some dipterans (Markow 2013; Murdy and Carson 1959; Stalker 1954; Stalker 1956), moths (Seiler 1960; Suomalainen *et al.* 1987), crustaceans (Nougue *et al.* 2015), and nematodes (Van der beek *et al.* 1998). Another mechanism of automictic parthenogenesis is terminal fusion. Here, the egg pronucleus fuses with its sister nucleus in the second-division polar body to form the zygote. With this mechanism, offspring from heterozygous mothers will become homozygous for either allele in the absence of recombination, but may retain maternal heterozygosity when there is recombination between locus and centromere. Terminal fusion automixis has been reported for example in mayflies (Sekine and Tojo 2010), termites (Matsuura *et al.* 2004), and orobatid mites (Heethoff *et al.* 2009). (Note however that in mites with terminal fusion automixis, meiosis may be inverted so that the consequences are the same as for central fusion (Wrensch *et al.* 1994).) Terminal fusion also seems to be the only confirmed mechanism of facultative parthenogenesis in vertebrates (reviewed in Lampert 2008). The most extreme mechanism of automixis is gamete duplication. Here, the egg undergoes either a round of chromosome replication without nuclear division or a mitosis followed by fusion of the resulting two nuclei. In both cases, the result is a diploid zygote that is completely homozygous at all loci. Gamete duplication has been reported in several groups of arthropods and in particular is frequently induced by inherited bacteria (*Wolbachia*) in hymenopterans (Gottlieb *et al.* 2002; Pannebakker *et al.* 2004; Stouthamer and Kazmer 1994). Finally, there are a number of genetic systems that are cytologically distinct from automixis but genetically equivalent or similar (see Discussion).

The peculiar mechanisms of automixis raise a number of questions. At the most basic level, one could ask why automixis exists at all. If there is selection for asexual reproduction, why not simply skip meiosis and produce offspring that are identical to their mother? Are there any advantages to automixis compared to clonal reproduction, or are there mechanistic constraints that make it difficult to produce eggs mitotically? How common is automixis, and how can it be detected and distinguished from other modes of reproduction using population genetic methods? What is the eventual fate of populations reproducing via automixis? Are the usual long-term problems faced by populations that forego genetic mixing (such as Muller’s ratchet or clonal interference) compounded in automictic populations because they also suffer from a form of inbreeding depression due to the a perpetual loss of heterozygosity, or can the loss of heterozygosity also be beneficial in some circumstances?

Answering these questions requires a firm understanding of how key evolutionary forces such as selection and drift operate in automictic populations. Although a few studies have yielded important insights into this issue, these studies have been limited to specific settings, e.g. dealing with the initial fitness of automictic mutants (Archetti 2004), selective maintenance of heterozygosity (Goudie *et al.* 2012), or the ‘contagious’ generation of new automictic lineages in the face of conflicts with the sex determination mechanism (EngelstÄDTER *et al.* 2011). Here, I develop mathematical models and report a number of analytical and numerical results on the evolutionary genetics in automictic populations. As a foundation, I first (re-)derive the expected distribution of offspring genotypes with up to two loci and under different modes of automixis in relation to crossover frequencies. Next, results on several statistics describing neutral genetic diversity are derived. I then investigate how natural selection on deleterious, overdominant and beneficial mutations operates in automictic populations. Finally, using the previous results I investigate the evolution of recombination suppression in automicts, a process that in extreme cases might effectively turn automictic into clonal reproduction.

## Models and Results

### Recombination and loss of heterozygosity

For a single locus, the relationship between crossover rates and loss of heterozygosity during automixis has been previously analyzed by several authors (EngelstÄDTER *et al.* 2011; Pearcy *et al.* 2006; Pearcy *et al.* 2011), so only a brief summary will be given here. The process can be understood by considering two steps. First, crossover events between the focal locus and its associated centromere during prophase I may or may not produce ‘recombinant’ genotypic configurations following meiosis I prophase (Fig. 1A). Second, the resulting configuration (Fig. 1B) may then either retain the original heterozygous state or be converted into a homozygous state (Fig. 1C).

In step 1, crossovers induce switches between two possible configurations (Fig. 1A). A crossover invariably produces a transition from the original state where sister chromatids carry the same allele to the state where sister chromatids carry different alleles, but only produces the reverse transition with probability 1/2. Based on this, it can be shown that with *n* crossovers, the probability of arriving in the ‘recombinant’ state is

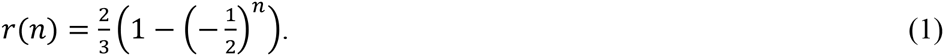

If we assume a Poisson distribution of crossover events with a mean of 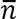, we obtain

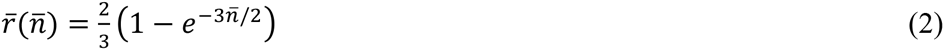

as the expected fraction of recombinant configurations. Of course, more complex distributions that take crossover interference into account could also be applied to Eq. (1) (Svendsen *et al.* 2015). It can be seen from Eqs. (1) and (2) that when the number of crossovers increases (i.e., with increasing distance of the locus from the centromere), the expected fraction of recombinant configurations converges to 2/3.

In step 2, the meiotic products form diploid cells through different mechanisms, resulting in three possible genotypes in proportions as shown in Fig. 1C. Combining Eq. (3) with these probabilities yields the following overall probabilities of conversion from heterozygosity in the mother to homozygosity in her offspring for central fusion (CF), terminal fusion (TF) and gamete duplication (GD) automixis:

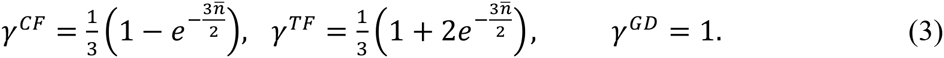

These equations can also be expressed in terms of map distance *d* (in Morgans) between the locus and its centromere that may be known in sexual conspecifics or related sexual species. This is done simply by replacing 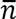 with *2d.*

Let us next consider two loci. Offspring proportions when at most one locus is heterozygous can be readily deduced from the single-locus case outlined above. Similarly, when the two loci are on different chromosomes, predicting offspring proportions is relatively straightforward because homozygosity will be attained independently at the two loci. When the two loci are on the same chromosome, predicting offspring proportions is more complicated. Now there are not two but seven distinct genotypic configurations following prophase of meiosis I, with transitions between these states induced by crossovers between the two loci and between the loci and their centromere (Figure S1). In the Supplementary Information (SI, section 1), the expected proportions of these configurations are derived both for fixed numbers of crossovers and assuming again a Poisson distribution of crossovers. Table 1 lists the corresponding proportions of genotypic configurations in a dually heterozygous mother for a number of scenarios of either complete linkage or absence of linkage. Also shown in Table 1 are the proportions of offspring genotypes resulting from each genotypic configuration under central fusion, terminal fusion and gamete duplication automixis. Finally, averaging over all genotypic configurations yields the total expected offspring distribution under the different automixis and linkage scenarios, as shown again for extreme linkage scenarios and the three modes of automixis in Table 2.

**Table 1:**
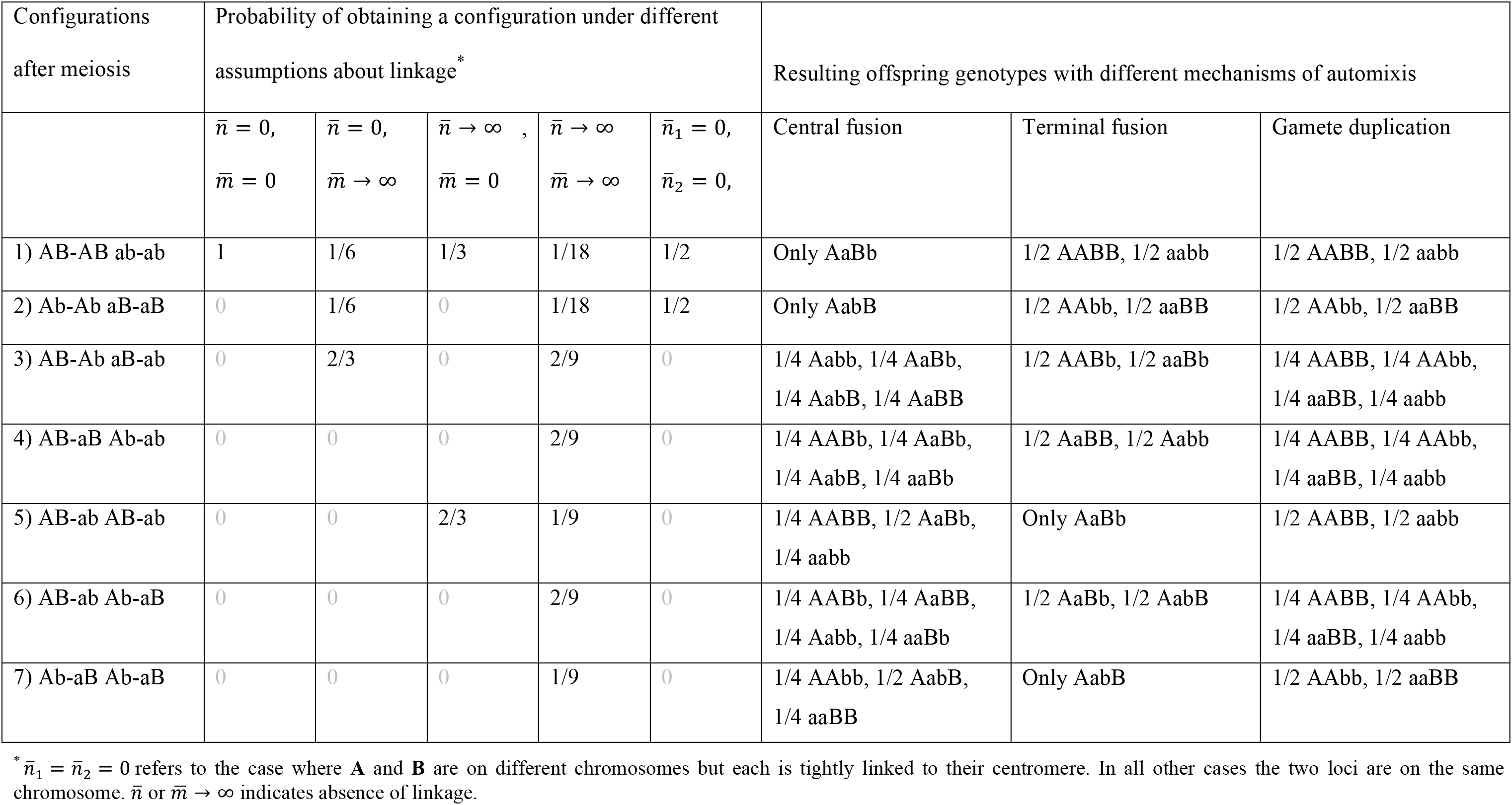
Postmeiotic genotype configurations and offspring genotypes arising from an *AaBb* mother.

**Table 2:** Total offspring distributions produced by AaBb mother.*

|  |  | AABB | AABb | AAbb | AaBB | AaBb | AabB | Aabb | aaBB | aaBb | aabb |
| --- | --- | --- | --- | --- | --- | --- | --- | --- | --- | --- | --- |
| Central fusion | $\bar{n} = 0, \bar{m} = 0$ | 0 | 0 | 0 | 0 | 1 | 0 | 0 | 0 | 0 | 0 |
| | $\bar{n} = 0, \bar{m} \rightarrow \infty$ | 0 | 0 | 0 | 1/6 | 1/3 | 1/3 | 1/6 | 0 | 0 | 0 |
| | $\bar{n} \rightarrow \infty, \bar{m} = 0$ | 1/6 | 0 | 0 | 0 | 2/3 | 0 | 0 | 0 | 0 | 1/6 |
| | $\bar{n} \rightarrow \infty, \bar{m} \rightarrow \infty$ | 1/36 | 1/9 | 1/36 | 1/9 | 2/9 | 2/9 | 1/9 | 1/36 | 1/9 | 1/36 |
| | $\bar{n}_1 = 0, \bar{n}_2 = 0$ | 0 | 0 | 0 | 0 | 1/2 | 1/2 | 0 | 0 | 0 | 0 |
| Terminal fusion | $\bar{n} = 0, \bar{m} = 0$ | 1/2 | 0 | 0 | 0 | 0 | 0 | 0 | 0 | 0 | 1/2 |
| | $\bar{n} = 0, \bar{m} \rightarrow \infty$ | 1/12 | 1/3 | 1/12 | 0 | 0 | 0 | 0 | 1/12 | 1/3 | 1/12 |
| | $\bar{n} \rightarrow \infty, \bar{m} = 0$ | 1/6 | 0 | 0 | 0 | 2/3 | 0 | 0 | 0 | 0 | 1/6 |
| | $\bar{n} \rightarrow \infty, \bar{m} \rightarrow \infty$ | 1/36 | 1/9 | 1/36 | 1/9 | 2/9 | 2/9 | 1/9 | 1/36 | 1/9 | 1/36 |
| | $\bar{n}_1 = 0, \bar{n}_2 = 0$ | 1/4 | 0 | 1/4 | 0 | 0 | 0 | 0 | 1/4 | 0 | 1/4 |
| Gamete duplication | $\bar{n} = 0, \bar{m} = 0$ | 1/2 | 0 | 0 | 0 | 0 | 0 | 0 | 0 | 0 | 1/2 |
| | $\bar{n} = 0, \bar{m} \rightarrow \infty$ | 1/4 | 0 | 1/4 | 0 | 0 | 0 | 0 | 1/4 | 0 | 1/4 |
| | $\bar{n} \rightarrow \infty, \bar{m} = 0$ | 1/2 | 0 | 0 | 0 | 0 | 0 | 0 | 0 | 0 | 1/2 |
| | $\bar{n} \rightarrow \infty, \bar{m} \rightarrow \infty$ | 1/4 | 0 | 1/4 | 0 | 0 | 0 | 0 | 1/4 | 0 | 1/4 |
| | $\bar{n}_1 = 0, \bar{n}_2 = 0$ | 1/4 | 0 | 1/4 | 0 | 0 | 0 | 0 | 1/4 | 0 | 1/4 |

\* $\bar{n}_1 = \bar{n}_2 = 0$ refers to the case where **A** and **B** are on different chromosomes but each is tightly linked to their centromere. In all other cases the two loci are on the same chromosome. $\bar{n}$ or $\bar{m} \rightarrow \infty$ indicates absence of linkage.

A general result from this analysis is that, as expected, the total fraction of offspring that become homozygous at each locus is the same as predicted by the single-locus equations. Thus, in the case of central fusion automixis and denoting by **A** the locus that is more closely linked to the centromere than the other locus **B**, we have

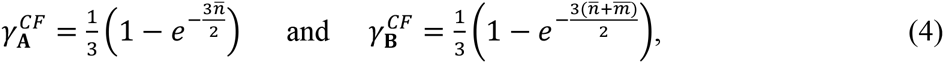

where 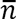 is the expected number of crossovers between the centromere and locus **A** and 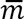 the expected number of crossovers between loci **A** and **B**. However, the two rates at which homozygosity is attained are not independent. Instead, since the two loci are linked, the fraction of offspring that are homozygous at both loci is greater than what is expected for each locus individually:

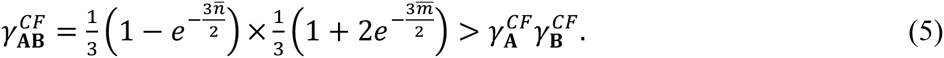

With terminal fusion, we obtain

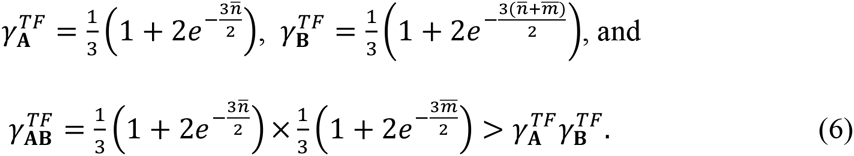

### Neutral genetic variation

Let us assume an unstructured, finite population of females reproducing through automictic parthenogenesis. We consider a single locus at which new genetic variants that are selectively neutral can arise by mutation at a rate μ and we assume that each mutation produces a new allele (the “infinite alleles-model”). Heterozygosity in individuals may be lost through automixis (with probability γ), and genetic variation can be lost through drift (determined by population size *N*). We are interested in the equilibrium level of genetic variation that is expected in such a population.

A first quantity of interest is the heterozygosity *H*_I_ i.e. the probability that the two alleles in a randomly chosen female are different. Following the standard approach for these type of models (e.g., Hartl and Clark 1997), the change in *H*_I_ from one generation to the next can be expressed through the change in homozygosity, 1 – *H*_I_ as

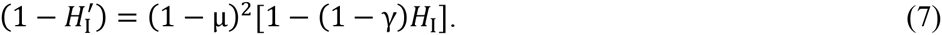

Solving 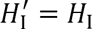 yields the equilibrium heterozygosity

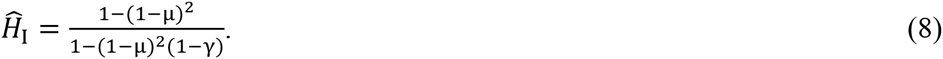

Second, we can calculate the probability *H*_T_ that two alleles drawn randomly from two different females are different (the “population level heterozygosity”). The recursion equation for *H*_T_ is given by

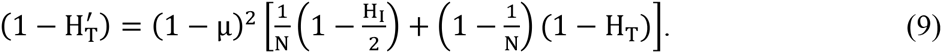

After substituting 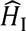 from Eq. (8) for *H*_I_, the equilibrium is found to be

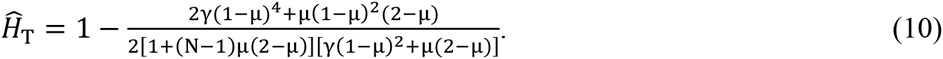

Combined, these two quantities allow us to calculate 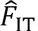, the relative difference between population-level and individual-level heterozygosity at equilibrium:

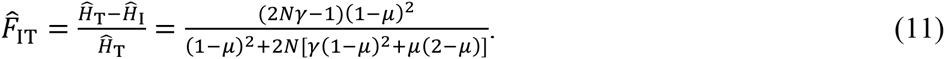

Finally, we can calculate the probability 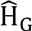 that two randomly drawn individuals have a different genotype at the locus under consideration. Unfortunately, this seems to require a more elaborate approach than the simple recursion equations used to calculate 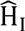 and 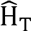, which is detailed in the SI (section 2). The resulting formula is rather cumbersome and not given here, but we can approximate 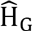 assuming *N*γ ≫ 1 and *Nμ* ≫ 1. This yields the (still unwieldy)

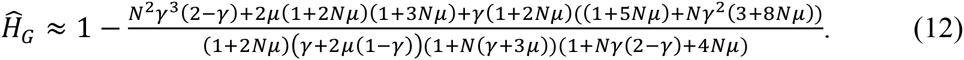

Figure 2 shows how the four statistics quantifying different aspects of genetic diversity depend on the rate γ at which heterozygosity in individuals is eroded, and compares them to the corresponding statistics in outbreeding sexual populations. Also shown in Figure 2 are diversity estimates from simulations (see SI, section 3 for details), indicating that the analytical predictions are very accurate. It can be seen that both the within-individual and population-level heterozygosities decline with increasing γ. Also, the former is greater than the latter for small values of γ, resulting in negative values of 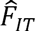, but this pattern reverses for larger γ. In Figure S2, the same relationships are shown but expressed in terms of map distance for the case of central fusion automixis (see Eq. 3). These plots illustrate that for the parameters assumed, large between-locus variation in equilibrium genetic diversity is only expected in the close vicinity of the centromeres.

**Figure 2.**
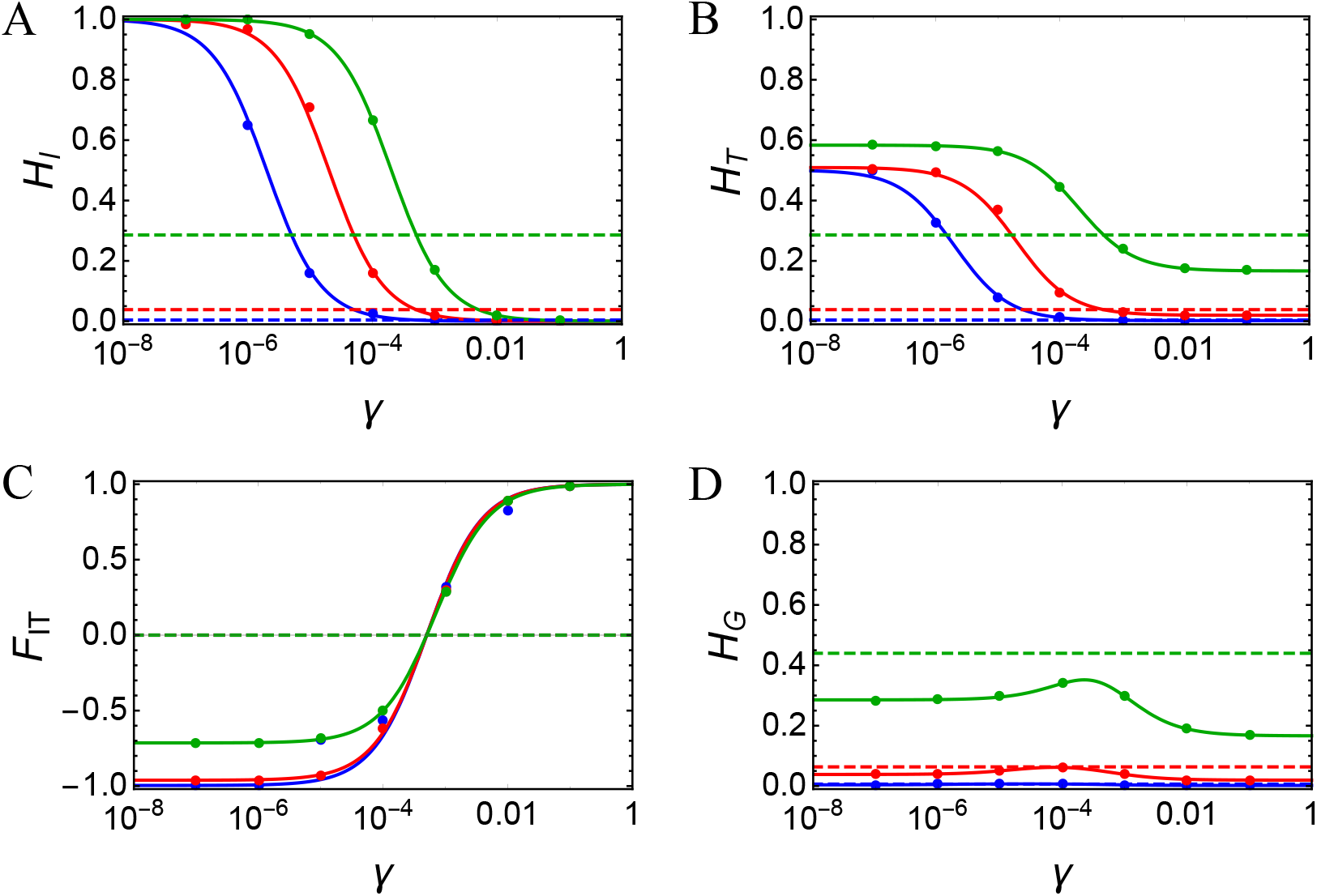
Equilibrium genetic diversity in neutrally evolving automictic populations, measured as A) within-individual heterozygosity 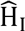, B) population-level heterozygosity 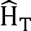, C) relative reduction in heterozygosity 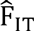, and D) diploid genotype-level diversity 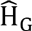. The solid lines in each plot show the analytical predictions for these statistics for varying rates γ at which heterozygosity is lost in the population, and for three mutation rates: 10^-7^ (blue), 10 (red) and 10 (green). Circles show the corresponding estimates from the simulations, and dashed lines show the corresponding expected values in outbreeding, sexual populations. Throughout, the population size was fixed to *N* = 1000. Note that most of the range of values for **y** shown are relevant only for central fusion automixis (0 **≤** *γ* **≤** 1/3), whilst predictions for terminal fusion and gamete duplications are restricted to the rightmost part of the plots (1/3 ≤ *γ* ≤ 1 and *γ* = 1, respectively).

In order to gain further insight into the formulae derived above, it may be helpful to consider a few special cases.

#### Special case 1: γ = 0

This case corresponds to strict clonal reproduction and has been studied previously (Balloux *et al.* 2003). In line with these previous results, all individuals are expected to eventually become heterozygous 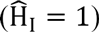, representing an extreme case of the Meselson effect (Normark *et al.* 2003; Welch and Meselson 2000). Furthermore, for the equilibrium population-level heterozygosity we get

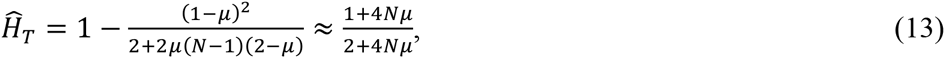

which is always greater than the corresponding genetic diversity in outbreeding sexual populations 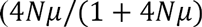, see also Fig. 2B). When 4*Nμ* is small, 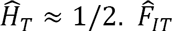 simplifies to

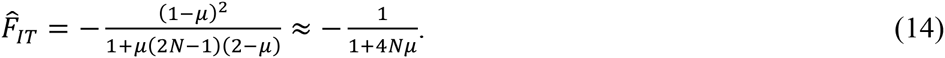

This is always negative because all individuals are heterozygous but alleles sampled from different individuals may still be identical. Finally,

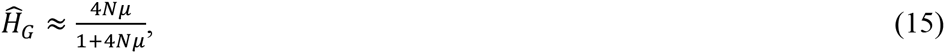

i.e. 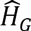 is identical to the equilibrium heterozygosity in sexual populations. This is because with strict clonal reproduction, each mutation gives rise to not only a new allele but also to a new genotype. New diploid genotypes arise twice as often as new alleles in sexual populations, but this is exactly offset by twice the number of gene copies in sexual populations than genotypes in clonal populations.

#### Special case 2: γ = 1 (gamete duplication)

This represents the opposite extreme: barring new mutations all heterozygosity is immediately lost. Thus, 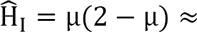 0 and, as a direct consequence, 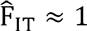. Moreover,

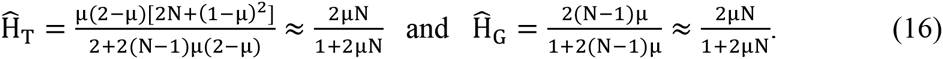

Not surprisingly, 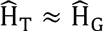 because when all individuals are homozygous, comparing sampled alleles from different females and comparing genotype samples is equivalent. Also, 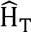 and 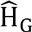 are always lower than the expected heterozygosity in sexual populations.

#### Special case 3: N → ∞

As the equilibrium heterozygosity is not affected by population size, equation (8) remains valid for 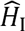. As expected both 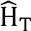 and 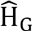 converge to one as *N* goes to infinity. Finally,

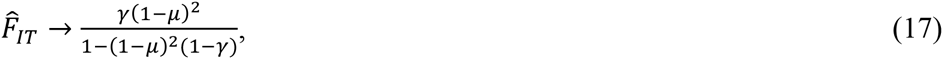

which is equal to the equilibrium individual-level homozygosity, 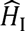.

### Mutation-selection balance

In order to investigate the balance between the creation of deleterious alleles through mutation and their purging by natural selection in automictic populations, let us assume an infinitely large population and a single locus with two alleles a (wildtype) and A (deleterious mutation). Wildtype aa individuals have a fitness of w_aa_ = 1 relative to heterozygotes with w_Aa_ = 1 – hs and mutant homozygotes with w_AA_ = 1 – s. To keep the model tractable, I assume that the mutation rate μ is small so that at most one mutation event occurs during reproduction, and that there is no back mutation from mutant to wildtype allele. Automixis operates as in the previous section, with heterozygotes producing a fraction γ/2 of either homozygote. Assuming a life-history order of selection-mutation-automixis, the recursion frequencies for the two mutant genotypes can be expressed as

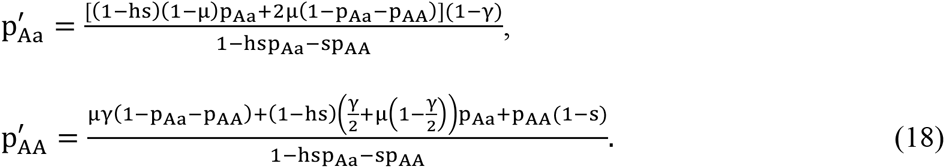

Solving 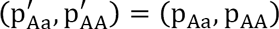 yields the frequencies of the mutant genotypes under mutation-selection equilibrium. Unfortunately, the resulting formulae are rather lengthy and uninformative. However, in a few special cases tractable results can be obtained.

#### Special case 1: Clonal reproduction (γ = 0)

In this case, the equilibrium that will be attained depends on the magnitude of the mutation rate. First, when the mutation rate is small, μ ≤ hs/(1 + hs), we get

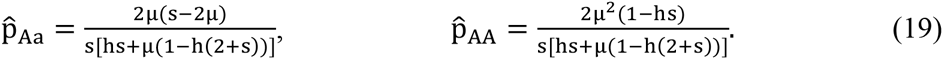

At this equilibrium, the population will consist mostly of mutation-free aa individuals, with some heterozygotes and very few AA homozygotes. Second, when hs/(1 + hs) < μ ≤ s/2, the equilibrium is given by

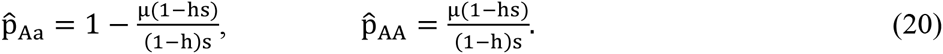

No mutation-free aa individual persist in the population in this case (since 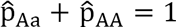). Intuitively, this situation arises when selection against heterozygotes is so weak relative to the mutation rate that eventually all aa individuals are converted into heterozygotes and a mutation-selection balance is attained between Aa and AA individuals. This balance is then analogous to the standard mutation-selection balance in haploid populations. Indeed, after re-normalizing all fitness values with the fitness of heterozygotes and defining an adjusted selection coefficient against AA homozygotes, 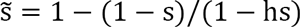, the equilibrium frequency of homozygous mutants can be expressed as 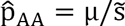. Finally, when μ > s/2, the mutation pressure outweighs selection completely and the mutant homozygotes will become fixed in the population 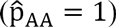.

#### Special case 2: Gamete duplication (γ = 1)

At the opposite extreme, when all heterozygotes are immediately converted into homozygotes, the equilibrium frequencies are given by

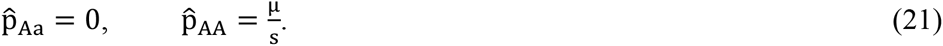

It is clear that in this case, selection against heterozygotes and thus the dominance coefficient *h* is irrelevant. Mutation-free aa homozygotes produce heterozygote mutant offspring at a rate 2µ per generation, but these heterozygotes are immediately converted into aa and AA offspring, each with probability 1/2. Thus, the effective rate at which AA offspring are produced is μ and the attained mutation-selection balance is identical to the one attained in haploid populations.

#### Special case 3: Recessive deleterious mutations (*h* = 0)

With arbitrary values of γ but strictly recessive deleterious mutations, the equilibrium is given by

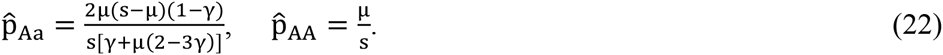

Thus, the equilibrium frequency 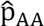 of mutant homozygotes is identical to the one expected in sexual populations with recessive deleterious mutations (corresponding to an equilibrium allele frequency of 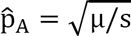). The equilibrium frequency of heterozygotes is a decreasing function of γ and can be either higher or lower than the corresponding equilibrium frequency of heterozygotes in sexual populations.

Figure 3 shows equilibrium frequencies of the Aa and AA genotypes under mutation-selection balance for recessive, partially recessive, semidominant and dominant mutations and compares these frequencies to the corresponding frequencies in sexual populations. For γ > 0, the frequency of heterozygotes is generally lower than in sexual populations whereas the frequency of AA homozygotes is higher than in sexual populations. Related to this, the equilibrium frequencies in automictic populations are generally much less sensitive to the dominance coefficient *h* than in sexual populations because γ > 0 implies that selection against heterozygotes is much less important than in sexual populations.

**Figure 3.**
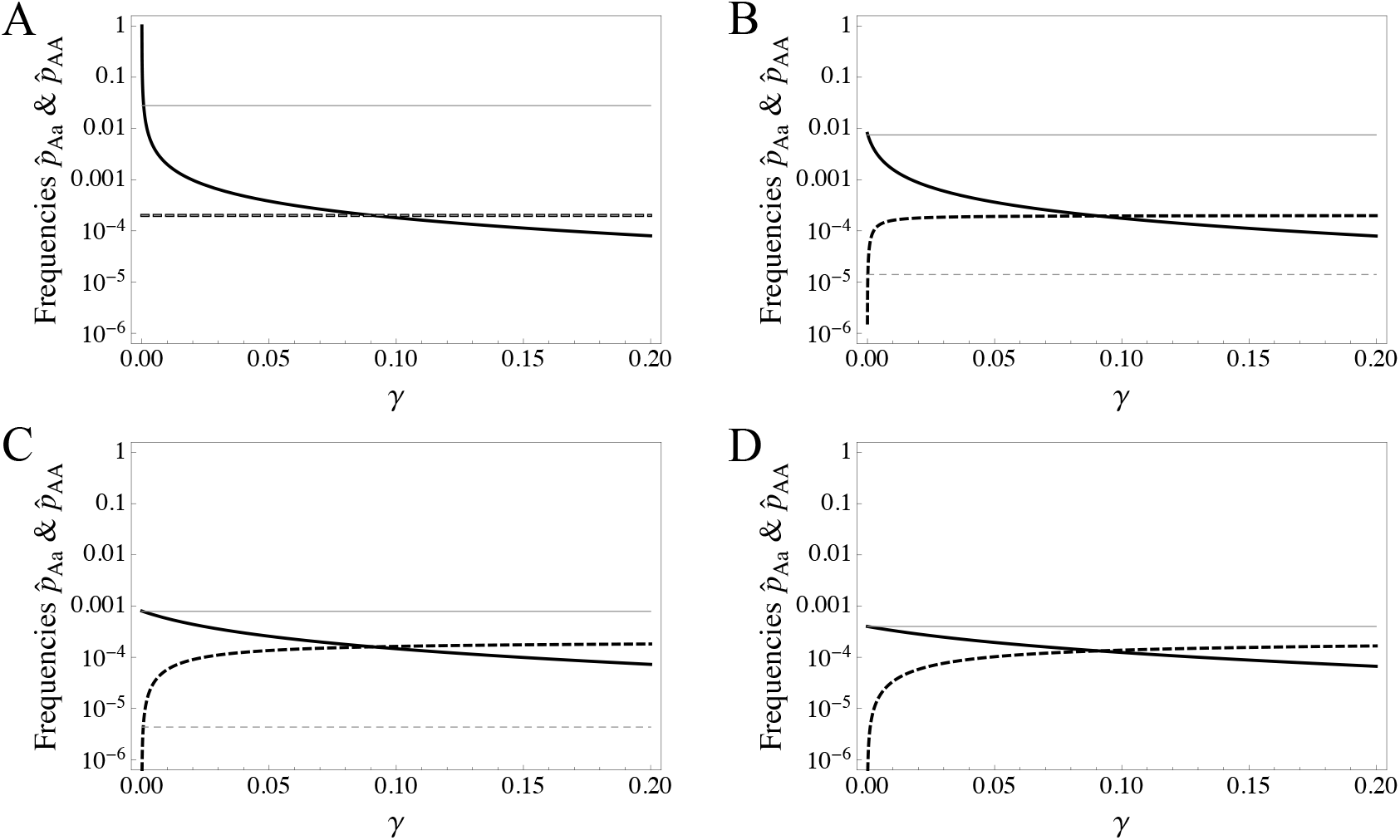
Equilibrium genotype frequencies under mutation-selection balance with different homozygosity acquisition rates γ, and for mutations that are A) recessive (*h* = 0), B) partially recessive (*h* = 0.05), C) semidominant (*h* = 0.5), and D) dominant mutations (*h* = 1). In each plot, the bold solid line gives the equilibrium frequency of *Aa* heterozygotes and the bold dashed line the equilibrium frequency of *AA* homozygotes. For comparison, the thin lines show the corresponding frequencies in outbreeding sexual populations. Other parameters take the values *s* = 0.05 and *μ* = 10^-5^.

The equilibrium frequencies can also be used to calculate the mutational load L_mut_, i.e. the relative reduction in the mean fitness of the population caused by recurrent mutation. This quantity can be expressed as 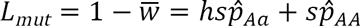. From this and Eqs. (20) and (21) it can be deduced that both with gamete duplication (γ = 1) and recessive mutations (*h* = 0), the genetic load in the population is given by the mutation rate μ, the same as for recessive mutations in sexual diploid populations and also the same as in haploid populations. In general, *L*_*mut*_ will be greater than μ, but, always slower than 2μ and, interestingly, often lower than the genetic load in sexual populations (Figure S3). This is in contrast to previous theoretical studies on the mutational load in clonal diploids in which the maintenance of heterozygosity caused the asexual populations to accumulate a higher load than the sexual populations (Chasnov 2000; Haag and Roze 2007). It is important to note however that the simple results derived here do not account for finite population size and interference between multiple loci, which may have a strong impact on mutation-selection balance and the mutational load (Glemin 2003; Haag and Roze 2007; Roze 2015).

### Overdominance

When there is overdominance (i.e., a heterozygote fitness advantage over the homozygote genotypes), it is useful to parameterize the fitness values as w_aa_ = 1 – s_aa_, w_Aa_ = 1 and w_AA_ = 1 **—** s_AA_, with 0 < s_aa_, s_AA_ ≤ 1. The recursion equation for the genotype frequencies p_Aa_ and p_AA_ can then be expressed as

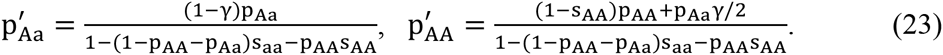

Solving 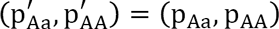 yields the following polymorphic equilibrium:

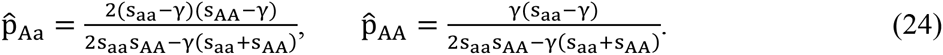

This equilibrium takes positive values for γ < s_aa_ and y < s_AA_, and stability analysis indicates that this is also the condition for the equilibrium to be stable. (The eigenvalues of the associated Jacobian matrix are (1 — s_aa_)/(1 — γ) and (1 — s_AA_)/(1 – γ).) Thus, overdominant selection can maintain heterozygotes in the face of erosion by automixis if the selection coefficient against either of the homozygotes is greater than the rate at which heterozygosity is lost. This result has previously been conjectured by Goudie *et al.* (2012) on the basis of numerical results of a similar model, and more complex expressions for equilibrium (24) have been derived by Asher (1970).

Depending on γ, the equilibrium frequency of heterozygotes can take any value between 0 (when γ ≥ min {s_aa_, s_AA_}) and 1 (when γ = 0, i.e. with clonal reproduction). This is shown in Figure 4 and contrasted with the equilibrium frequency in sexual populations, given by 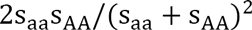.

**Figure 4.**
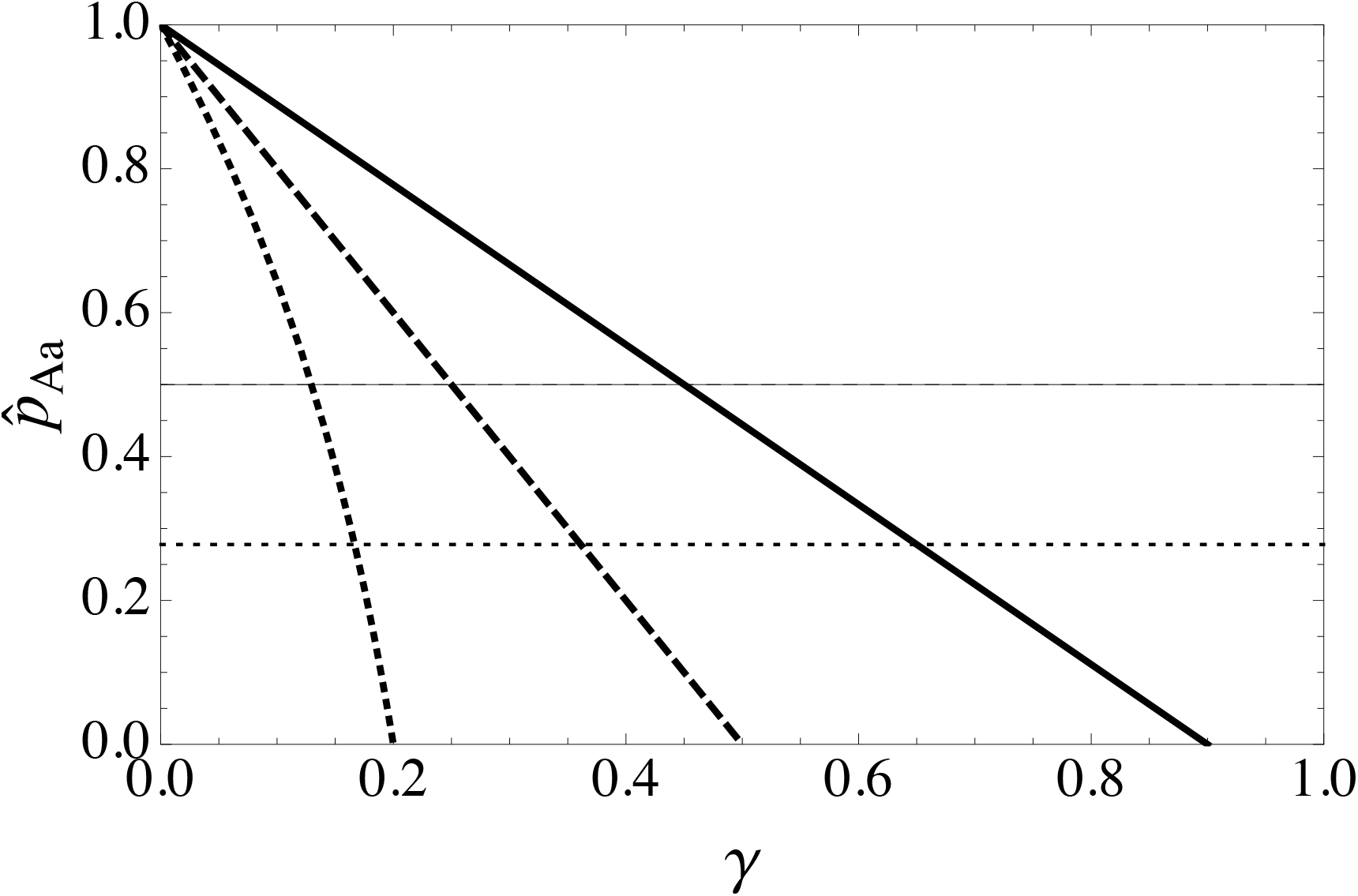
Equilibrium heterozygote frequencies under overdominant selection and for given rates *γ* at which heterozygotes are converted into homozygotes. The bold lines show these frequencies under automictic reproduction and for *s*_*AA*_ = *s*_*aa*_ = 0.9 (solid line), *s*_*AA*_ = *s*_*aa*_ = 0.5 (dashed line), and *s*_*AA*_ = 1, *s*_*aa*_ = 0.2 (dotted line). For comparison, the thin lines show the corresponding frequencies in outbreeding sexual populations.

We can also calculate how much the mean fitness in the population at equilibrium is reduced by automixis compared to clonal reproduction. Provided that 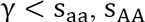, this “automixis load” is given by

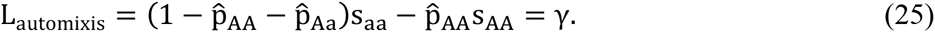

This simple formula parallels the classic result that the mutational load in haploid populations is given by the mutation rate and thus shows that automixis acts like mutation in producing two genotypes (AA and aa) of inferior fitness from the fittest genotype (Aa) that are then purged by natural selection. The genetic load can be either smaller or greater than the corresponding segregation load in a sexual population. More precisely, when 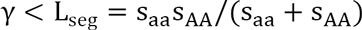 (Crow and Kimura 1970), there will be more heterozygotes in the automictic population and their genetic load will be lower than in the sexual population, and *vice versa.* When γ > min {s_aa_, s_AA_}, the heterozygotes are lost from the population and either the aa (if *s*_*aa*_ < *s*_*AA*_) or the AA genotype (if *s*_*aa*_ > *s*_*AA*_) will become fixed. In this case, we obtain the largest possible load, 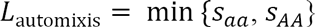.

### Associative overdominance

In addition to overdominance, heterozygosity could also be maintained through off-phase recessive deleterious mutations at tightly linked loci (Frydenberg 1963; Ohta 1971), and this has been proposed to explain heterozygosity in the Cape honeybee (Goudie *et al.* 2014). Consider a recently arisen lineage reproducing through central fusion automixis in which by chance, the founding female carries a strongly deleterious recessive mutation *A* on one chromosome and another strongly deleterious recessive mutation *B* at a tightly linked locus on the homologous chromosome. Thus, the genetic constitution of this female is *AabB.* Then, the vast majority of offspring that have become homozygous for the high-fitness allele *a* are also homozygous for the deleterious allele *B* and *vice versa*, so that linkage produces strong indirect selection against both *aa* and *bb* homozygotes.

In order to explore this mechanism, numerical explorations of a two-locus model of an infinitely large population undergoing selection and reproduction through automixis were performed. This model builds upon the expressions for loss of heterozygosity in the presence of recombination between a centromer and two loci derived above (for details see SI section 4). Heterozygosity at either locus entails a reduction of fitness of 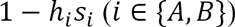 whereas homozygosity for the deleterious alleles reduces fitness by 1 − *s*_*i*_ in each locus. Fitness effects at the two loci are multiplicative (i.e., no epistasis). An example run is shown in Figure 5A. As can be seen, the *AabB* genotype is maintained at a high frequency for a considerable number of generations (in automixis-selection balance with the two homozygous genotypes *AAbb* and *aaBB*) before it is eroded by recombination between the two loci and the *aabb* genotype spreads to fixation. We can also treat the two linked loci as a single locus and use the equilibrium frequency of heterozygotes derived above for single-locus overdominance to estimate the quasi-stable frequency of the *AabB* genotype before it is dissolved. Specifically, this frequency can be approximated by Eq. (24) following substitution of *S*_*AA*_ for *S*_*B*_, *S*_*aa*_ for *S*_*A*_, and *γ* for the expression in Eq. (3), yielding

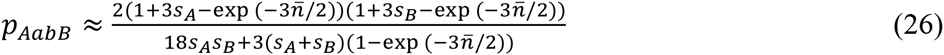

**Figure 5.**
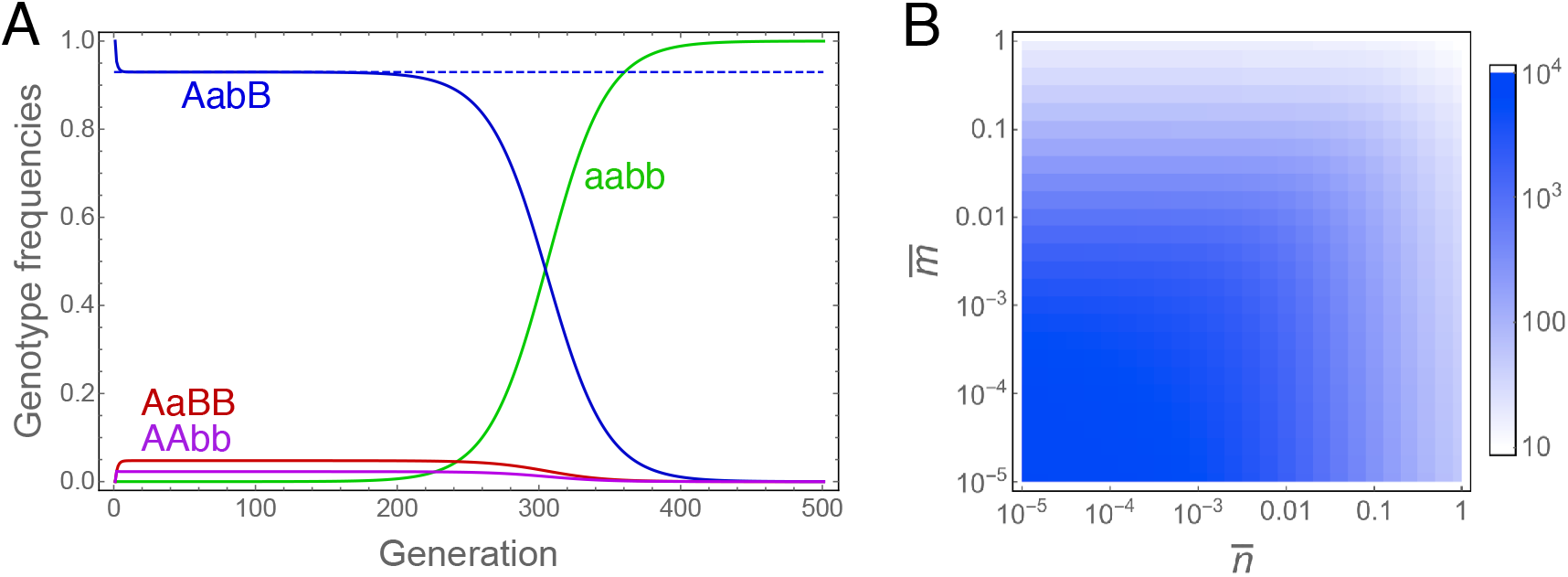
Maintenance of strongly deleterious mutations (A and B) in an *AabB* genotype through associative overdominance with central fusion automixis. A) Example evolutionary dynamics where solid lines show the four predominant genotypes in the population and the dashed blue line gives the approximation for the quasi-stable frequency of the *AabB* genotype. Parameters take the values *s*_*A*_ = 0.999, *s_B_ =* 0.5, *h_A_ = h_B_ =* 0, 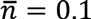 and 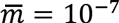 = 10^-7^. B) Time until dissolution of the *AabB* genotype for different mean crossover numbers 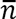 (between centromer and locus **A**) and 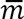 (between loci **A** and **B**). Values shown are the number of generations until the frequency of the *AabB* genotype has dropped from 1 to 0.01 in the population. Other parameters take the values *s_A_ = s_B_ =* 0.99 and *h_A_ = h_B_ =* 0.001.

For simplicity, this approximation assumes complete recessivity (*h*_*A*_ = *h*_*B*_ = 0), but partial recessivity could also readily be incorporated. As shown in Fig. 5A, this approximation is very close to the quasi-stable frequency of the *AabB* genotype obtained numerically.

Next, we can ask for how long the *AabB* genotype is expected to persist in the population. To address this question, screens of the parameter space with respect to the two mean crossover numbers were performed. The recursion equations were initiated with only *AabB* individuals present in the population and iterated until their frequency dropped below 0.01. The number of generations this took for different mean numbers of crossovers 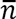 between locus **A** and the centromere and different numbers of crossovers 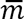 between loci **A** and **B** are shown in Figure 5B. It can be seen that provided that the two loci are both tightly linked to the centromer, central fusion automixis can indeed maintain the polymorphism for many generations. The same principle also applies to terminal fusion automixis and sexual reproduction, but here the deleterious recessive mutations are only maintained for very short time periods (results not shown).

### Spread of beneficial mutations

In order to better understand adaptive evolution in automictic populations, consider first a deterministic single locus model without mutation and with relative fitness 1 + *hs* and 1 + *s* for heterozygotes and *AA* homozygotes, respectively. The recursion equations for this model can then be written as

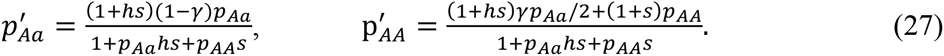

Assuming that both the *Aa* and the *AA* genotype are rare in the population and that *s* is small, these recursion equations can be approximated by

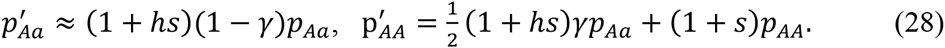

This system of recursion equations can be solved, and if we further assume that initially there are only one or few heterozygote mutants but no AA homozygotes (*p*_*AA*_(0) = 0), this solution becomes

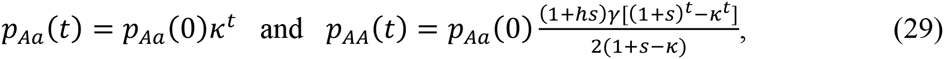

with 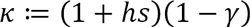. These expressions demonstrate that when *γ* is large relative to the selection benefit of heterozygotes — more precisely when *κ* < 1, or 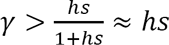 – the heterozygotes will not be maintained and the beneficial mutation will instead spread as a homozygous genotype through the population. Thus, in this case we have for sufficiently large *t*:

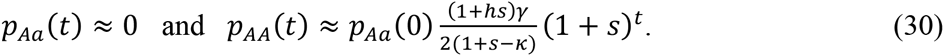

Here, *h* and *γ* determine how efficiently the initial heterozygotes are maintained and converted to homozygotes, but only *s* determines the actual rate at which the beneficial mutation spreads. By comparison, a beneficial mutation in an outbreeding sexual population will initially be found in heterozygotes only, with

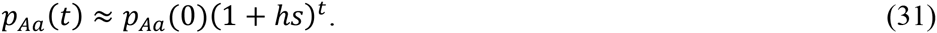

Thus, the rate at which the mutation spreads in sexual populations is determined by the fitness advantage in heterozygotes only, which means the mutation will always spread at a lower rate than in automictic populations. Nevertheless, the heterozygotes in sexual populations have a ‘head start’ relative to the homozygotes in automictic populations (see fraction in Eq. 30), which results from the fact that only half of the original heterozygotes are converted into homozygotes. This means with high dominance levels *h,* it might still take some time until a beneficial allele reaches a higher frequency in an automictic compared to a sexual population.

When 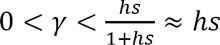, both heterozygotes and AA homozygotes will spread simultaneously in the population and, for very small *γ*, it may take a long time until the homozygotes reach a higher frequency than the heterozygotes. In the extreme case of *γ* = 0 (clonal reproduction), heterozygote frequency increases by a factor of (1 + *hs*) in each generation (i.e., at the same rate as in outbreeding sexuals) and no homozygotes are produced.

These considerations clearly show that unless beneficial mutations are completely dominant, they will spread faster in automictic than in either clonal or sexual populations. To what extent does this result hold when more than one locus is considered? It is well known that asexually reproducing populations may fix beneficial mutations more slowly because of clonal interference, i.e. competition between simultaneously spreading beneficial mutations that in the absence of recombination cannot be brought together into the same genome (Fisher 1930; Muller 1932). Despite the term “clonal interference”, this mechanism should also operate in automictic populations, and we can ask whether and under what conditions the decelerating effect of clonal interference on the speed of adaptive evolution can offset the beneficial effects of turning heterozygous with one beneficial into homozygotes with two beneficial mutations.

To address this question, numerical investigations of a two-locus model involving recombination, automixis, selection and drift were performed. (Note that clonal interference only operates in finite populations subject to random genetic drift and/or random mutations; full details of this model can be found in the SI, section 5.) The results of a screen of the two crossover rates 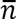 (between locus **A** and the centromere) and crossovers 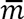 between (between loci **A** and **B**) are shown in Fig. 6. It appears that unless 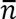 is very small, recessive beneficial mutations spread considerably faster in automictic than in sexual populations, despite clonal interference in the former. With additive fitness effects, the difference between automictic and sexual populations is less pronounced and the beneficial effects of recombination become apparent when 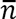 is very small. Perhaps surprisingly, the mean number 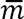 of crossover between the two loci under selection, which determines the recombination rate in the sexual population, plays only a minor role and needs to take low values for recessive beneficial mutations to spread slightly faster in sexual than automictic populations (bottom-left corner in Fig. 6A). This is because although increasing values of 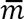 lead to faster spread of the beneficial mutations in sexual populations, this effect is only weak compared to the accelerating effect of increasing 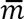 in automictic populations.

**Figure 6.**
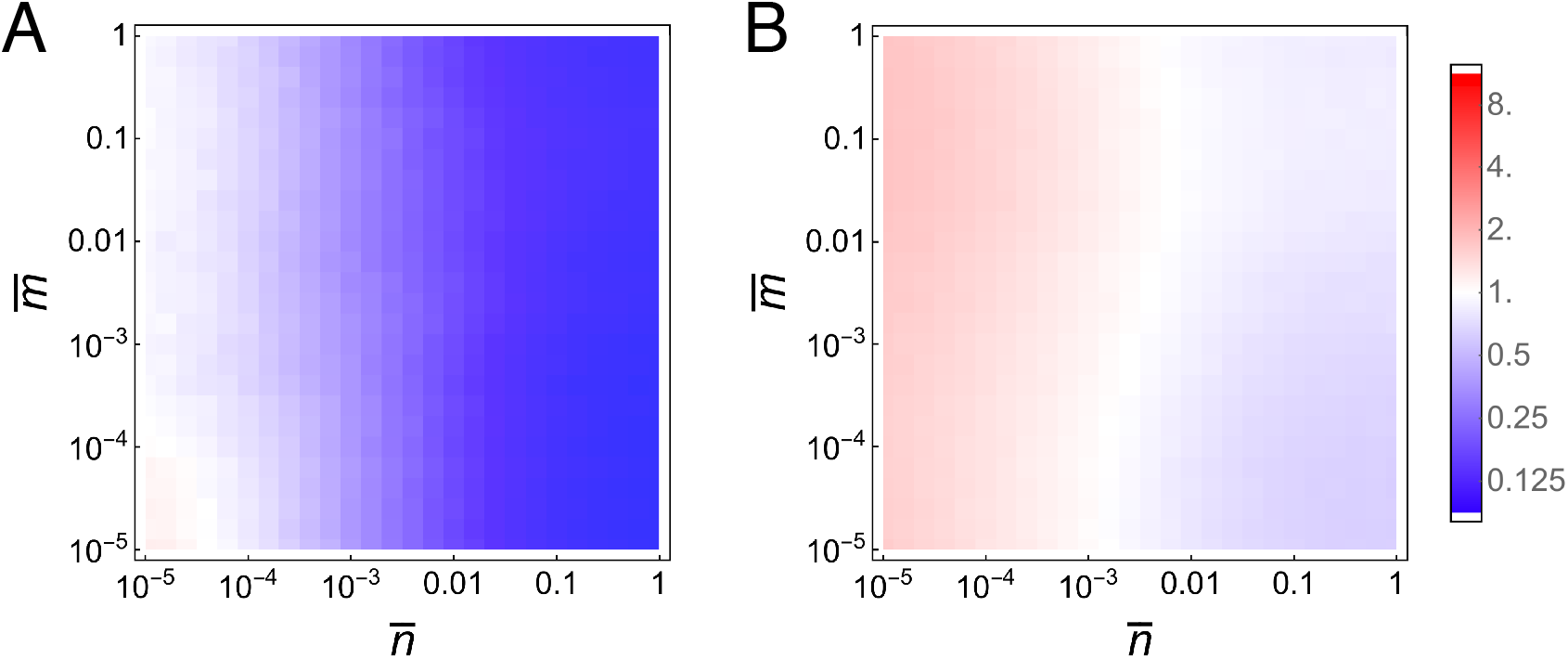
Number of generations required for the spread of two beneficial mutations in populations reproducing through central fusion automixis relative to sexual populations. Each plot shows these relative times for different mean crossover numbers 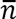 (between the centromere and locus **A)** and 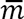 (between loci **A** and **B**). Blue color (relative times < 1) indicates that the spread of the beneficial mutations was faster in the automictic than in the sexual populations, and red color (relative times >1) indicates the opposite. Each simulation was initialized with a population of non-adapted *aabb* genotypes. Beneficial mutations *A* and *B* could then arise at a rate μ = 10^-5^ and were assumed to provide a fitness benefit of *s*_*A*_ = *s*_*B*_ = 0.1, with dominance coefficient A) *h*_*A*_ = *h*_*B*_ = 0.01 (recessive beneficial mutations) or B) *h*_*A*_ = *h*_*B*_ = 0.5 (additive effects). Populations were subject to random genetic drift with population size *N*=10,000 and results were averaged over 1000 replicate simulations. Spread of the beneficial mutations was considered complete when the mean population fitness had increased by more than 99% of the maximum possible increase.

### Selection on crossover rates

We finally turn to the question of whether natural selection is expected to reduce crossover rates in automictic populations. Let us first consider a population reproducing by central fusion automixis in which heterozygosity at a given locus is maintained through overdominant selection. Combining the results from Eq. (3) and (25), the mean fitness of a resident population with a mean crossover number 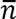 is given by

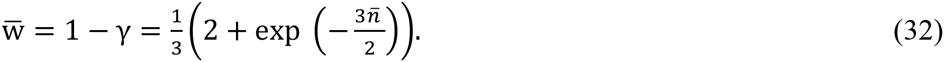

Since reproduction is asexual and assuming a dominant crossover modifier allele, the selection coefficient σ for a mutant genotype with a different crossover rate can be obtained simply by comparing mean fitness of the resident and the mutant lineage. (This is in contrast to recombination rate evolution in sexual populations, where a much more sophisticated approach is required (Barton 1995).) If the factor by which crossovers numbers are altered is denoted by *α* (i.e., the mutant mean number of crossovers is 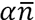), this yields

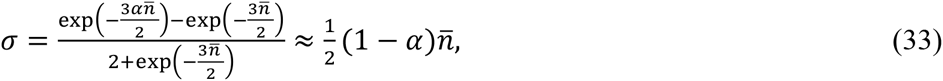

where the approximation is valid for small 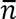. As expected, any mutant in which crossover rates are suppressed (*α* < 1) is selectively favoured (σ > 0). In the extreme case of *α* = 0 (complete crossover suppression),

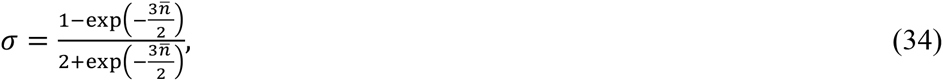

which ranges from around 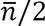 when the initial crossover rate is already very small to 1/2 when 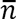 is very large.

With terminal fusion automixis, we obtain

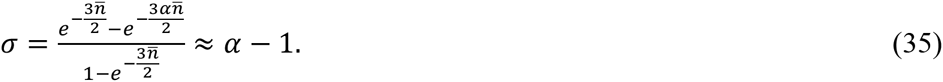

Again, the approximation is valid for small 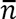 (but note that conditions where overdominance stably maintains heterozygosity are rather limited in this case; see Eq. (24)). Increases in crossover rates are selected for with terminal fusion, but even when there are many crossovers between the focal locus and its centromere, a substantial genetic load (*L* = 1/3) will persist.

We can also ask how selection should operate on crossover rates in automictic populations evolving under mutation-selection balance. Without answering this question in any quantitative detail, we can note that since the equilibrium genetic load decreases with increasing *γ* (Fig. S3), there should be selection for increased crossover rates in populations with central fusion automixis and selection for decreased recombination rates in populations with terminal fusion automixis. However, given that genetic load is always in the range between *μ* and 2*μ*, selection for increased crossovers will be only very weak on a per-locus basis (*σ* < *μ*).

Preliminary numerical investigations competing two lineages with different crossover rates confirm the predictions on selection on crossover rates. However, it will be important to study this problem more thoroughly in a multi-locus model and with finite populations so that for example also the impact of stochastically arising associative overdominance can be ascertained.

## Discussion

### Automixis as a viable system of reproduction?

Automixis is a peculiar mode of reproduction. Not only are, as with other modes of asexual reproduction, the benefits of recombination forfeited, but the fusion of meiotic products to restore diploidy also means that heterozygosity can be lost at a high rate. This raises intriguing questions as to why automixis has evolved numerous times and how stably automictically reproducing populations can persist. Previous work has shown that the loss of complementation faced by a newly evolved automictically reproducing female can cause severe reductions in fitness that may exceed the twofold cost of sex and are likely to severely constrain the rate at which sex is abandoned (Archetti 2004; Archetti 2010; EngelstÄDTER 2008). However, the results obtained here indicate that once an automictic population is established, it may persist and in some respects even be superior to clonal or sexual populations. In particular, neutral genetic diversity will be lower in automictic than in clonal populations but may still be greater than in sexual populations, the mutational load will generally be lower in automictic than in both sexual and clonal populations (unless mutations are completely recessive), and the genetic load caused by overdominant selection can be lower in automictic than in sexual populations.

Empirical examples confirming that automicts can be highly successful at least on short to intermediate timescales include the Cape honeybee “Clone” (which has been spreading for more than 25 years) (Goudie and Oldroyd 2014), the invasive ant *Cerapachys biroi* (which has been reproducing asexually for at least 200 generations) (Oxley *et al.* 2014; Wetterer *et al.* 2012), and *Muscidifurax uniraptor* wasps which have been infected by parthenogenesis-inducing *Wolbachia* for long enough that male functions have degenerated (Gottlieb and Zchori-FEIN 2001). Of course, automictic populations still suffer from the lack of recombination and hence long-term consequences such as the accumulation of deleterious mutations through Muller’s ratchet or reduced rates of adaptation because of clonal interference. It therefore does not come as a surprise that like other asexuals, automictic species tend to be phylogenetically isolated (Schwander and Crespi 2009). One exception to this rule are the oribatid mites, in which around 10% out of >10,000 species reproduce by automixis and radiations of automictic species have occurred (Domes *et al.* 2007; Heethoff *et al.* 2009). In order to better understand the long-term dynamics of adaptation and mutation accumulation in automictic populations, it would be useful to develop more sophisticated models than presented here that incorporate multiple loci and random genetic drift.

### Relationship to other genetic systems

There is a bewildering diversity of genetic systems that have similarities to automixis. In order to discuss the relationship of the results obtained here with prior work it may be useful to group these genetic systems into two classes. The first are systems that are mechanistically distinct from automixis but are genetically equivalent. This includes systems in animals and plants where there is no fusion of meiotic products but some other meiotic modification that has the same consequences, and also systems of intratetrad mating. The results obtained here are thus directly applicable and previous theoretical work on such systems can directly be compared to the work presented here. For example, parthenogenesis in *Daphnia pulex* has been reported to proceed through a modified meiosis in which the first anaphase is aborted halfway, homologous chromosomes are re-joined and the second meiotic division proceeds normally (Hiruta *et al.* 2010). Similarly, some forms of apomixis in plants (meiotic diplospory) are also achieved by suppression of the first meiotic division (Gustaffson 1931; Van DIJK 2009). These modifications of meiosis are genetically equivalent to central fusion automixis and can, through complete suppression of recombination, also lead to clonal reproduction. Intratetrad mating is commonly found in many fungi, algae and other organisms and is achieved through a variety of mechanisms (Hood and Antonovics 2004; Kerrigan *et al.* 1993). Provided the mating-type locus is completely linked to the centromere, intra-tetrad mating is genetically equivalent to central fusion automixis (Antonovics and Abrams 2004). If the mating-type locus is not closely linked to the centromere, the outcome would still be equivalent to automixis but with a mixture of terminal and central fusion, depending on whether or not there has been a recombination event between the mating-type locus and the centromere.

The second class of systems comprises those that are very similar to automixis but equivalent only when a single locus is considered. This means that many of the results reported here (e.g., on neutral variation, mutation-selection balance and overdominance) can still be applied. For example, clonal populations in which there is occasional, symmetrical gene conversion can be considered genetically equivalent to the single-locus models considered here, with the rate of loss of heterozygosity *γ* replaced by the gene conversion rate. Gene conversion has been reported in several parthenogenetic animals (Crease and Lynch 1991; Flot *et al.* 2013; Schon and Martens 2003), and recently a number of results concerning coalescent times and patterns have been derived for such systems (Hartfield *et al.* 2016). Populations that reproduce exclusively by selfing also belong into this class; such populations are characterized by a rate of heterozygosity loss of *γ* = 1/2. It should be emphasized, however, that selfing in general is rather distinct even from random automixis: in essence, the difference is that alleles are sampled either with (selfing) or without (automixis) replacement from the meiotic products.

### Populations with mixed automictic and sexual reproduction

The models presented here assume populations that reproduce exclusively through automixis. Although several species with exclusively automictic reproduction have been reported, many other species exhibit different forms of mixed sexual and automictic reproduction. The simplest case is one where a lineage of automicts competes with sexual conspecifics but where there is no gene flow between these two populations. Such a situation is found in the Cape honeybee, *Apis mellifera capensis*, in which a subpopulation (the “Clone”) reproduces through central fusion automixis and parasitizes colonies of a sexual subspecies, *A. mellifera scutellata* (Goudie and Oldroyd 2014). In principle, the results presented here could be used to predict the outcome of such competitions by comparing population mean fitness of sexual and automictic populations. However, the case of the Cape honeybee is fraught with a number of additional complexities, including both honeybee characteristics such as eusociality and the complementary sex determination system, and the parasitic nature and the epidemiological dynamics of the Clone. This will make it necessary to develop specifically tailored models that incorporate both the evolutionary genetics consequences of automixis explored in the present paper and the ecological and genetic idiosyncrasies of the Clone (Martin *et al.* 2002; Moritz 2002).

More complicated is the case of gene flow between the sexual and automictic subpopulations. This can occur for example when otherwise automictic females occasionally produce males. Provided these males are viable and fertile, they may mate and produce offspring with the sexual females. This will not only introduce genetic material from the asexual into the sexual populations, but it may also lead to the emergence new automictic lineages because the males may transmit the genes coding for automictic reproduction to their female offspring. Such cases of ‘contagious parthenogenesis’ (Simon *et al.* 2003) associated with automixis have been reported in the parasitoid wasp *Lysiphlebus fabarum* (Sandrock *et al.* 2011; Sandrock and Vorburger 2011) and *Artemia* brine shrimps (Maccari *et al.* 2014). Some aspects of the evolutionary dynamics of such systems have been studied (EngelstÄDTER *et al.* 2011), but population genetic processes such as the ones studied here remain to be investigated.

Finally, there are many species in which there are no clear sexual and asexual subpopulations but where females can reproduce both sexually and through automixis. This includes for example the majority of *Drososophila* species where parthenogenesis has been reported (Markow 2013), and also a number of vertebrates with facultative parthenogenesis (Lampert 2008). It is expected that sexual populations capable of occasional automictic reproduction should not differ much from sexual populations in terms of population genetic processes. One exception is that rare automixis may facilitate the colonization of previously uninhabited areas. Distinguishing the genomic signature of the resulting automictically arisen population bottlenecks from those of ‘conventional’ bottlenecks will be challenging but may be feasible with data on genome-wide levels of heterozygosity (see also Svendsen *et al.* 2015). On the other hand, rare sex in predominantly automictic populations is expected to have a great impact as the mixing of lineages may efficiently counteract clonal interference and Muller’s ratchet (Hojsgaard and Horandl 2015).

### Conclusions

In this study, a number of theoretical results regarding basic on the population genetics of automictic populations were derived both for neutral and selective processes. A general conclusion that emerges is that in analogy to strong levels of inbreeding, automictic reproduction is difficult to evolve but once established may be viable on intermediate timescales and even has advantages compared to clonal and sexual reproduction. Future theoretical work is still necessary to elucidate long-term evolutionary patterns of automictic species such as the rate of mutational meltdown under Muller’s ratchet or the dynamics of adaptation.

## Acknowledgments

I thank Christoph R. Haag and Nicholas M. A. Smith for their insightful comments on the manuscript. I would also like to thank the Instituto Gulbenkian Ciência (Portugal) for hosting me during the final stages of preparing this manuscript.

---

**Figure S1.**
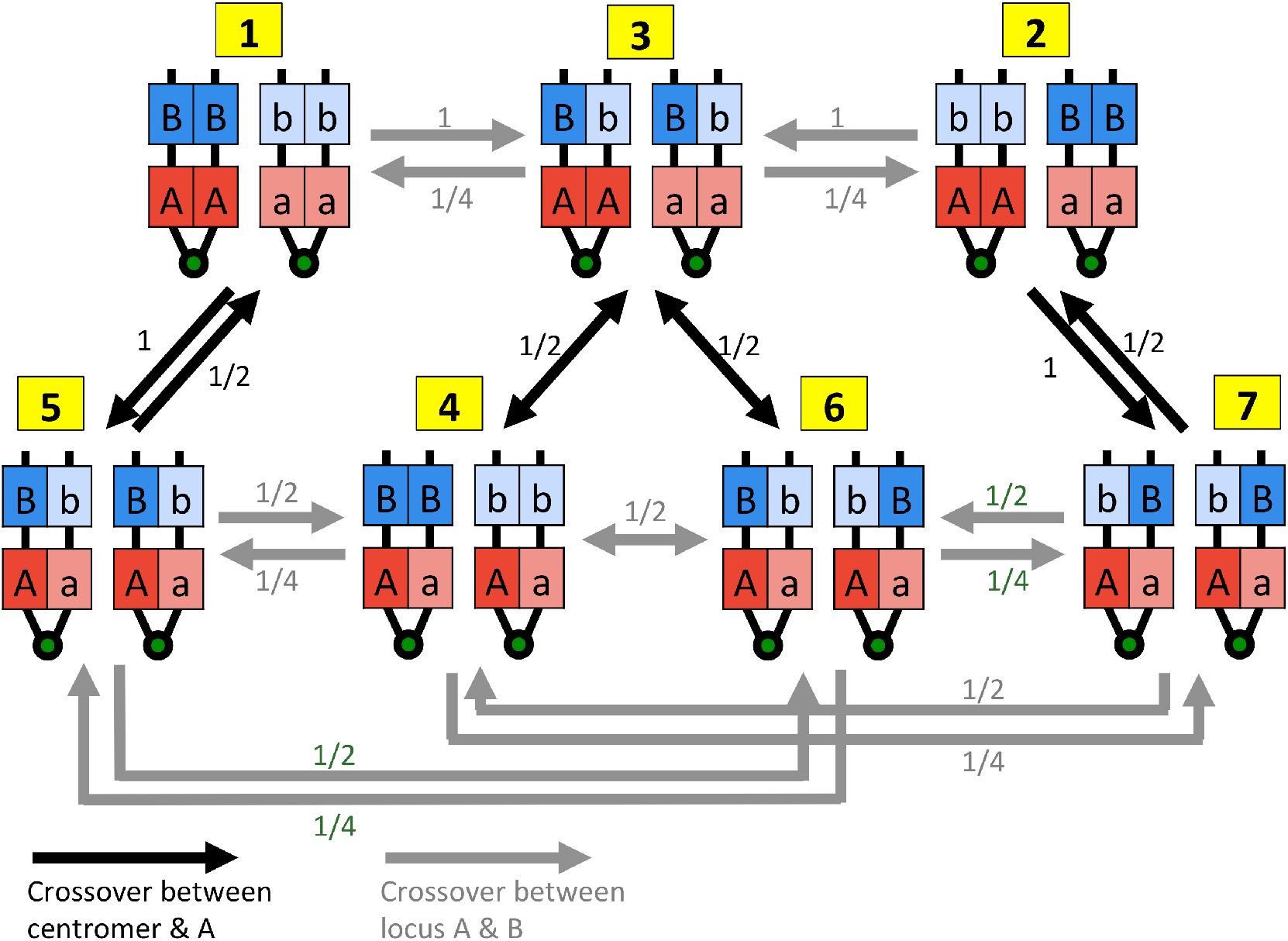
Illustration of the seven possible genotypic states with two linked loci following meiotic prophase I, and how crossovers between the two loci (grey) or between the locus **A** and the centromere (black) effect transitions between these states. Numbers next to the arrows indicate probabilities and the numbering of the states (yellow boxes) corresponds to the states as defined in the SI, section 1.

**Figure S2.**
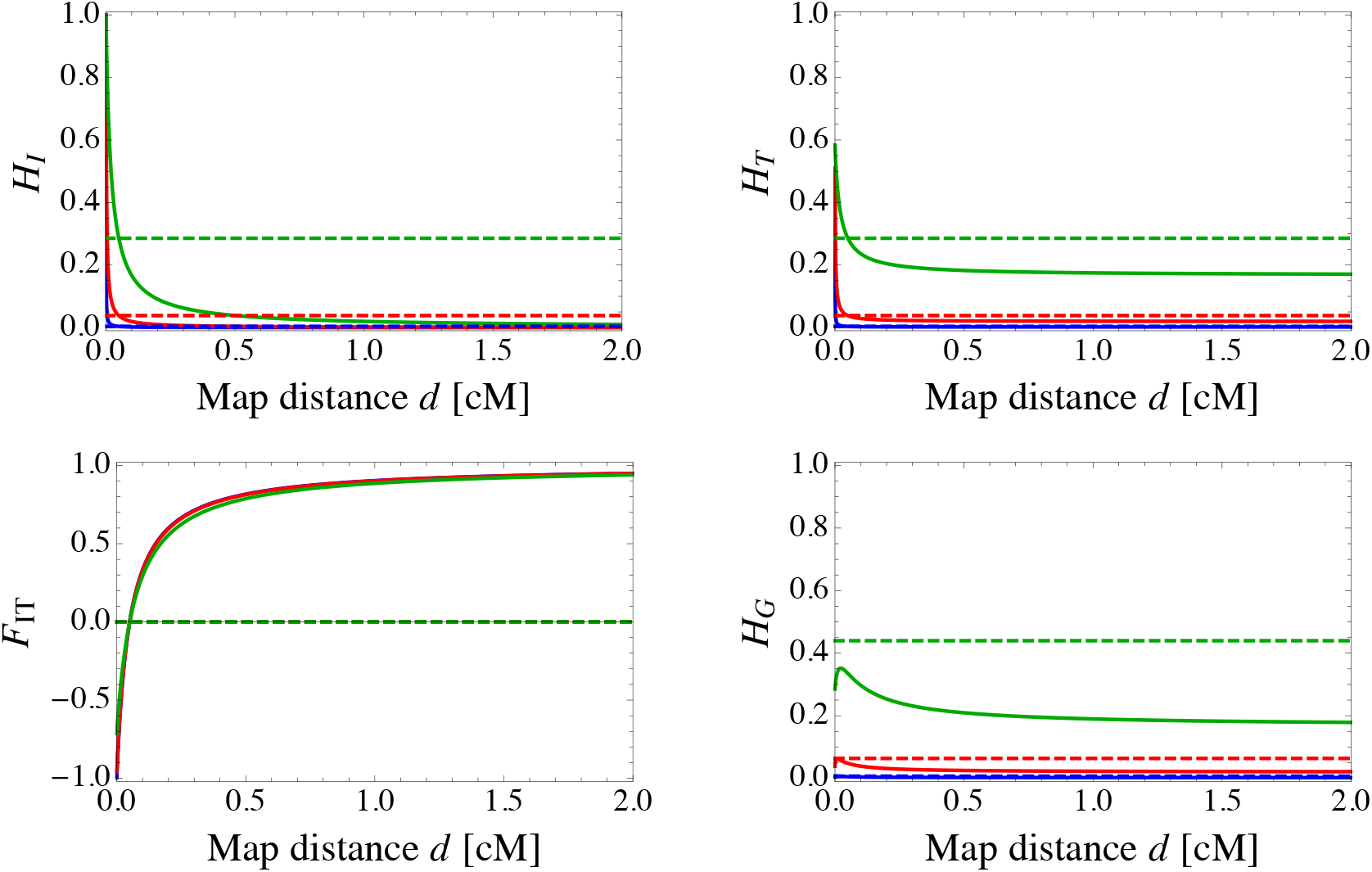
Equilibrium genetic diversity in neutrally evolving populations reproducing by central fusion automixis. The same analytical results as in Fig. 2 are shown and the same parameters are used (*N*=1000, μ=10^-7^ (blue), 10^-6^ (red) and 10^-5^ (green)), but the x-axis is re-scaled to show map distance in centimorgans.

**Figure S3.**
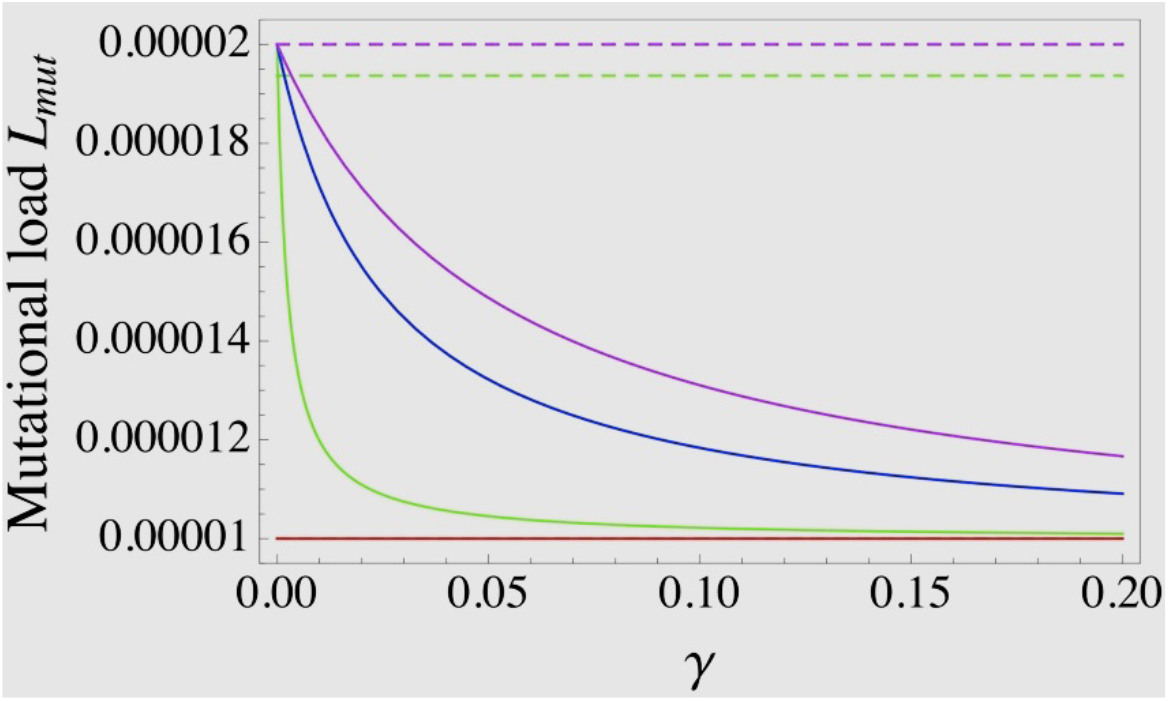
Mutational load under mutation-selection balance depending on the rate of heterozygosity degradation *γ*. Solid lines show mutational loads in automictic populations and dashed lines show the corresponding loads in outbreeding sexual populations. Parameters take the values *s* = 0.05, *μ* = 10^-5^ and *h* = 0 (red), *h* = 0.05 (green), *h* = 0.5 (blue), and *h* = 1 (purple). Note that for the latter two values of *h*, the mutational loads in sexual populations are indistinguishable in this plot, and that for completely recessive mutations, the genetic load in both sexual and automictic populations is equal to *μ*.

